# PARP-1, EpCAM, and FRα as potential targets for intraoperative detection and delineation of endometriosis: a quantitative tissue expression analysis

**DOI:** 10.1101/2023.12.19.572311

**Authors:** Beatrice Belmonte, Giovanni Di Lorenzo, Alessandro Mangogna, Barbara Bortot, Giorgio Bertolazzi, Selene Sammataro, Simona Merighi, Anna Martorana, Gabriella Zito, Federico Romano, Anna Giorgiutti, Cristina Bottin, Fabrizio Zanconati, Andrea Romano, Giuseppe Ricci, Stefania Biffi

## Abstract

Endometriosis is a gynecological disease characterized by the presence of endometrial tissue in abnormal locations, leading to severe symptoms, inflammation, pain, organ dysfunction, and infertility. Surgical removal of endometriosis lesions is crucial for improving pain and fertility outcomes, with the goal of complete lesion removal. Incomplete surgery often results in recurring symptoms and the need for additional interventions. This study aims to explore the biological significance of PARP-1, EpCAM, and FRα as potential targets for molecular imaging in endometriosis, assessing their suitability for targeted intraoperative imaging.

Gene expression analysis was performed using the EndometDB. By immunohistochemistry, we investigated the presence and distribution of PARP-1, EpCAM, and FRα in endometriosis foci and adjacent tissue. We also applied an ad hoc platform for the analysis image to perform a quantitative immunolocalization analysis. Additionally, double immunofluorescence analysis was carried out for PARP-1 and EpCAM, as well as PARP-1 and FRα, to explore the expression of these combined markers within endometriosis foci and their potential simultaneous utilization in surgical treatment.

The analysis of gene expression revealed that PARP-1, EpCAM, and FRα exhibit higher levels of expression in endometriotic lesions compared to the peritoneum, used for control tissue. Immunohistochemical analysis also highlighted a significant increase in expression of all three biomarkers in endometriosis foci compared to surrounding tissues, as well as the quantitative analysis of immunolocalization of the signal. Furthermore, the double immunofluorescence analysis consistently revealed nuclear expression of PARP-1 and membrane and cytoplasmatic expression of EpCAM and FRα, although relatively weaker.

Overall, the three markers demonstrate significant potential for effective imaging of endometriosis. Particularly, the results emphasize the importance of PARP-1 expression as a possible indicator for distinguishing endometriotic lesions from adjacent tissue. PARP-1, as a potential biomarker for endometriosis, offers promising avenues for further investigation in terms of both pathophysiology and diagnostic-therapeutic approaches.

## Introduction

Endometriosis is a debilitating gynecological condition defined by the implantation of endometrial tissue in ectopic places and usually associated with persistent inflammation resulting in pain, organ dysfunction, and infertility [1,2]. Endometriosis most commonly occurs in the lower abdomen or pelvis, but it can appear anywhere in the body [2,3]. Pelvic endometriosis can manifest as ovarian cysts (ovarian endometrioma), superficial peritoneal endometriosis (SPE), and deep infiltrating endometriosis (DIE), depending on the site and depth of implantation [1,4]. DIE involves nodular lesions which invade the surrounding organs beneath the peritoneum. They are more aggressive and commonly found on the uterosacral ligaments, bladder, vagina, and intestine [5]. The complete surgical removal of endometriosis lesions can improve both pain symptoms and fertility outcomes, with the primary therapy goal being the removal of all visible lesions [6]. On the other hand, incomplete surgery is frequently associated with the recurrence of pain symptoms and the need for repeat surgery, with significant associated morbidity. Therefore, improvements in the resection technique are highly sought to improve outcomes, decrease functional loss and recurrence, and increase the patient’s quality of life. Minimally invasive surgery is the preferred surgical approach because it is usually associated with less pain, shorter hospital stays, and faster recovery [7]. Although endoscopy allows for magnification of the operative field, the identification of endometriosis implants is not always possible using white light because some implants may be very small or hidden, especially in case of DIE covered by healthy peritoneum or small superficial implant extending deeply retroperitoneum. Studies report that enhanced imaging allows for detecting of additional endometriotic lesions missed by conventional white-light laparoscopy [7,8].

Fluorescent dyes such as fluorescein and indocyanine green (ICG) have been used for many years in clinical practice [9]. They can outline the vascular system and help identify areas of high perfusion or permeability. However, fluorescent dyes present drawbacks, such as speedy body clearance and the lack of precise targeting properties [10]. During the last years, molecular imaging probes have been developed to get around the limits of free dyes and visualize and measure biological activity in living systems [10,11]. A molecular imaging probe includes a signal agent and a targeting moiety [12,13]. The signal agent generates signals for imaging, whereas the targeting moiety interacts with a biomarker.

Intraoperative fluorescent molecular imaging is mainly used in cancer treatment, but it also has the potential to treat benign disorders like endometriosis [11]. Endometriosis is linked to gynecologic oncology, particularly ovarian cancer [14,15]. The tumor niche in ovarian cancer and the pro-endometriotic niche in endometriosis, in particular, exhibit considerable chronic inflammatory and immunosuppressive characteristics [14].

Epithelial cell adhesion molecule (EpCAM) was one of the first cancer-related biomarkers to be identified. The first monoclonal antibody, edrecolomab, being tested on patients over thirty years ago [16]. Recent findings suggest that overexpression of EpCAM, accompanied by an epithelial-mesenchymal transition (EMT), might be involved in endometriosis [17]. In a previous study, both EpCAM and folate receptor alpha (FRα) seemed to be promising for targeted intraoperative imaging of endometriosis [18]. FRα is found on the apical surface of epithelial cell membranes in endometriotic lesions but it is not found in the surrounding normal tissue [18]. Nowadays, clinical interest has focused on the role of poly (ADP-ribose) polymerase 1 (PARP-1) which could represent a potential target for molecular imaging probes and therapy. Increased PARP-1 activity has been linked to several cancers and inflammation-related pathologies, such as asthma, sepsis, arthritis, atherosclerosis, and neurodegenerative disorders [19]. It has been suggested that PARPis, designed for cancer therapy, might be potentially used for inflammatory disorders treatment [19].

The main goal of this study was to investigate the presence and distribution of PARP-1, EpCAM, and FRα to identify a potential target for molecular imaging in endometriosis. As these three markers appear to have promising potential for targeted intraoperative imaging of endometriosis, we investigated their expression as both intensity signal and topographic distribution within endometriosis foci, comparing them with surrounding tissues [18].

## Methods

### Patients’ cohort

Tissue samples were selected from a retrospective database created by the IRCCS Burlo Garofolo, comprising specimens from patients who had undergone surgery for endometriosis. Following approval by the Institutional Review Board (IRB-BURLO 01/2022, 09.02.2022), patients were asked to sign an informed consent form. Tissue samples from 11 consecutive patients were used in the present study.

The inclusion criteria for patients’ selection were the histological confirmation of superficial peritoneal endometriosis (SPE) and deep infiltrating endometriosis (DIE). Exclusion criteria: interdicted patients, unable to understand informed consent; patients with other forms of endometriosis without endometriosis reported in inclusion criteria; peritoneal inflammatory diseases (i.e., pelvic inflammatory disease, diverticulosis, etc.); oncological patients with peritoneal involvement.

### Gene expression analysis

We explored differentially expressed genes in peritoneum and endometriosis lesions using the EndometDB, freely accessible at https://endometdb.utu.fi/. The En-dometDB, a public database, includes the expression data from 115 patients and 53 controls, with over 24,000 genes and clinical characteristics, such as age, disease stages, hormonal medication, menstrual cycle phase, and endometriosis lesion types [20].

### Immunohistochemical analyses

For immunohistochemical (IHC) analysis, four-micrometer–thick formalin-fixed and paraffin-embedded (FFPE) tissue sections were deparaffinized, rehydrated, and unmasked using Epitope Retrieval Solutions (Novocastra) at pH6 and pH9 in thermostatic bath at 98°C for 30 minutes. Subsequently, slides were washed in PBS at room temperature. After endogenous peroxidase neutralization with 3% H_2_O_2_ and Fc blocking with 0.4% casein in PBS (Novocastra), sections were incubated with antibodies.

We used the following primary antibodies: rabbit anti-human PARP-1 (clone EPR18461, 1:100 pH6, Abcam), rabbit anti-human EpCAM (1:400 pH9, Abcam), rabbit anti-human FRα (1:1000 pH9, Thermofisher).

IHC staining was revealed using Novolink Polymer Detection System (Novocastra) and DAB (3,3’-Diaminobenzidine, Novocastra) as substrate chromogen.

Double fluorescent immunostainings for PARP-1/EpCAM and PARP-1/FRα were performed using Opal Multiplex IHC kit (Akoya Biosciences).

After deparaffinization, antigen unmasking was carried out with Epitope Retrieval Solution (pH9, Novocastra) boiled at 100% power, followed by 20% power for 15 min using microwave technology (MWT). Sections were incubated with Blocking Buffer for 10 min at room temperature and then with primary antibody for 90 min at room temperature. Slides were then incubated with Polymeric Horseradish Peroxidase-conjugated (HRP) secondary antibody for 10 min, and signal was developed using Opal 520 fluorophore-conjugated tyramide signal amplification (TSA, 1:100 dilution). To allow the next antigen detection, slides were again processed with microwave treatment for primary-secondary antibody complexes stripping; then they were incubated with second primary antibody, followed by Polymeric Horseradish Peroxidase-conjugated (HRP) secondary antibody and Opal 620 fluorophore-conjugated tyramide signal amplification (TSA, 1:100 dilution). Finally, sections were microwaved in Antigen Retrieval Buffer and nuclei were subsequently visualized with DAPI (4′,6-diamidino-2-fenilindole).

Slides were analyzed with Axioscope A1 microscope (Zeiss) equipped with four fluorescence channels widefield IF. Microphotographs were collected using Axiocam 503 color digital camera with Zen 2.0 Software (Zeiss).

### Quantitative immunolocalization analysis

Quantitative analyses of IHC staining were performed by calculating the average percentage of positive signals in endometriosis foci within all samples at low-power magnification (×100) using the Nuclear Hub (weak positivity: signal intensity threshold 210; moderate positivity: signal intensity threshold 188; strong positivity: signal intensity threshold 162) or Positive Pixel Count v9 (1+ weak positivity: signal intensity range 220–175; 2+ moderate positivity: signal intensity range 175–100; 3+ strong positivity: signal intensity range 100–0) ImageScope software.

### Statistical analysis

The paired two sample Bootstrap t-test [21] was applied to compare the average percentages between two groups (lesions vs surrounding tissues). The boot.t.test function from the R package MKinfer was used to calculate the p-values. The p-values have been adjusted using the Benjamini-Hochberg correction. The differences with adjusted p-value < 0.05 were considered to be significant.

## Results

### Patients’ selection and clinical data

Tissue samples from 11 patients were used in the present study. The patient’s characteristics and medication are reported in **Table 1**. Collected surgical samples, their anatomical location, and pathological description are reported in **Table 2**.

**Table 1.**
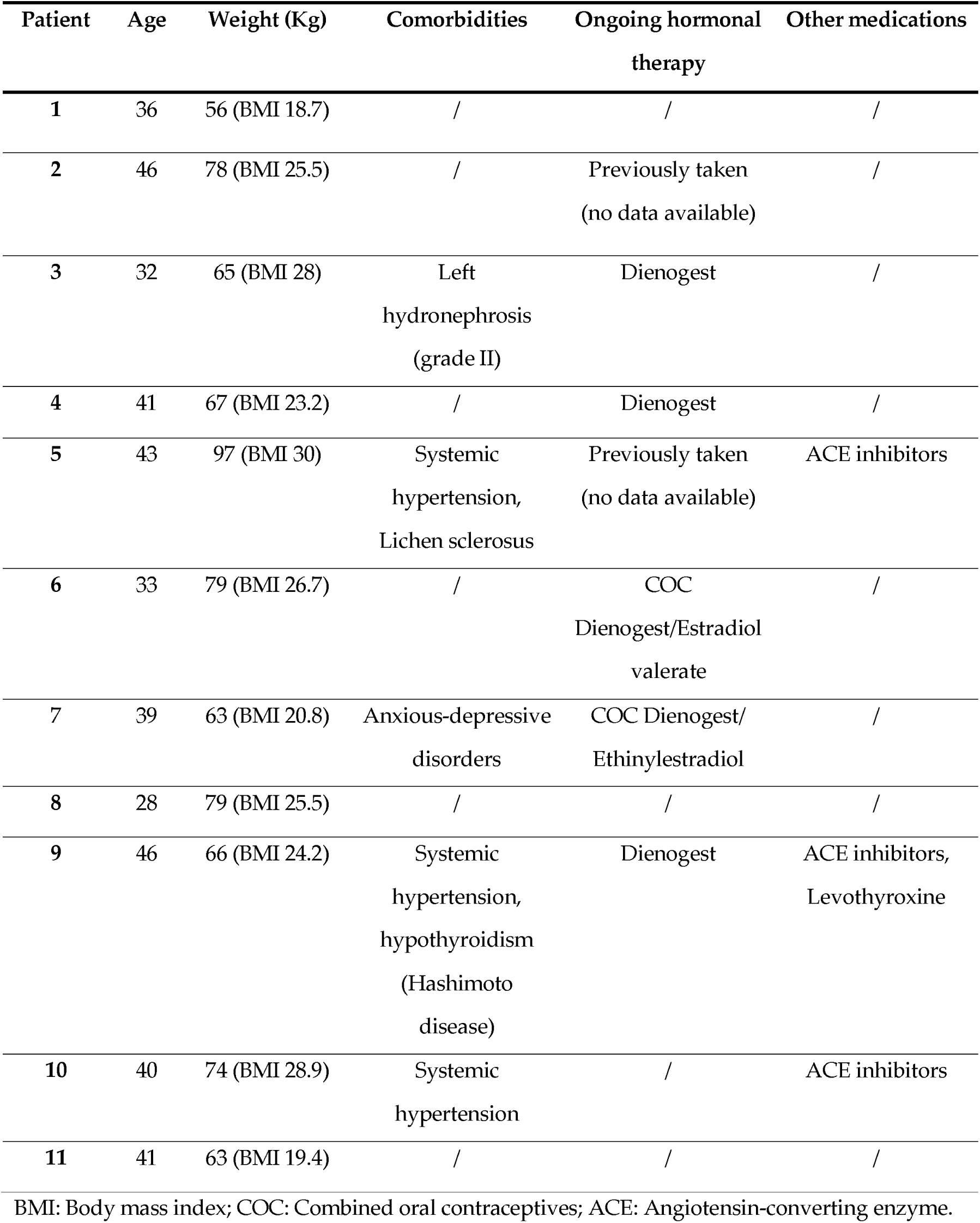
Patient’s characteristics.

**Table 2.**
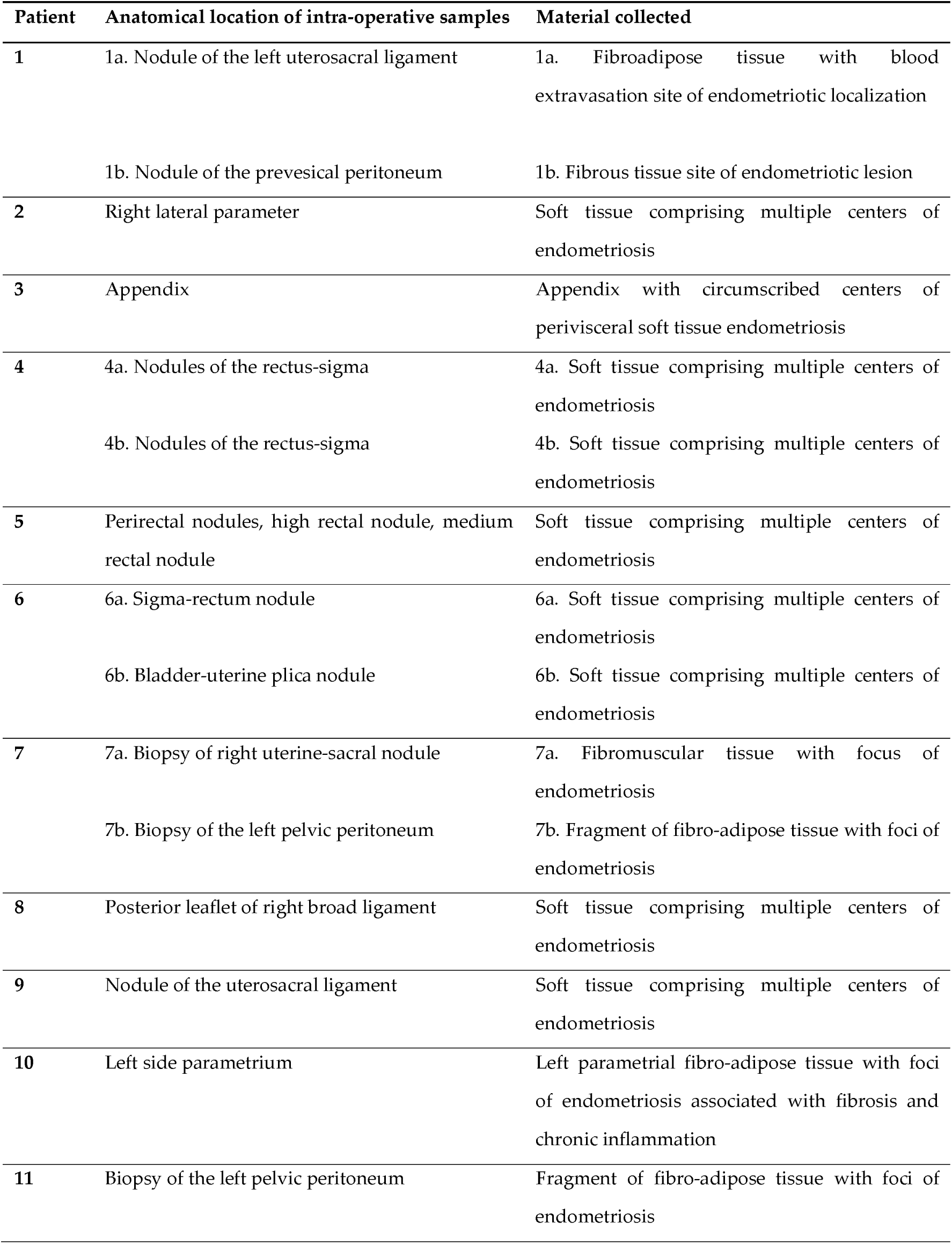
Surgical samples characteristics.

| **Patient** | **Anatomical location of intra-operative samples** | **Material collected** |
| --- | --- | --- |
| **1** | 1a. Nodule of the left uterosacral ligament | 1a. Fibroadipose tissue with blood extravasation site of endometriotic localization |
|  | 1b. Nodule of the prevesical peritoneum | 1b. Fibrous tissue site of endometriotic lesion |
| **2** | Right lateral parameter | Soft tissue comprising multiple centers of endometriosis |
| **3** | Appendix | Appendix with circumscribed centers of perivisceral soft tissue endometriosis |
| **4** | 4a. Nodules of the rectus-sigma | 4a. Soft tissue comprising multiple centers of endometriosis |
|  | 4b. Nodules of the rectus-sigma | 4b. Soft tissue comprising multiple centers of endometriosis |
| **5** | Perirectal nodules, high rectal nodule, medium rectal nodule | Soft tissue comprising multiple centers of endometriosis |
| **6** | 6a. Sigma-rectum nodule | 6a. Soft tissue comprising multiple centers of endometriosis |
|  | 6b. Bladder-uterine plica nodule | 6b. Soft tissue comprising multiple centers of endometriosis |
| **7** | 7a. Biopsy of right uterine-sacral nodule | 7a. Fibromuscular tissue with focus of endometriosis |
|  | 7b. Biopsy of the left pelvic peritoneum | 7b. Fragment of fibro-adipose tissue with foci of endometriosis |
| **8** | Posterior leaflet of right broad ligament | Soft tissue comprising multiple centers of endometriosis |
| **9** | Nodule of the uterosacral ligament | Soft tissue comprising multiple centers of endometriosis |
| **10** | Left side parametrium | Left parametrial fibro-adipose tissue with foci of endometriosis associated with fibrosis and chronic inflammation |
| **11** | Biopsy of the left pelvic peritoneum | Fragment of fibro-adipose tissue with foci of endometriosis |

### Gene expression analysis with EndometDB shows that PARP-1, EpCAM, and FRα are overexpressed in the endometriotic lesion compared to the peritoneum

We examined the expression levels of PARP-1, EpCAM, and FRα in the peritoneum and both peritoneal and deep lesions using data from the EndometDB database (**Figure 1**). All three markers demonstrate a significant increase in expression in lesions compared to peritoneal tissue. Deep lesions have a lower expression than peritoneal lesions.

**Figure 1.**
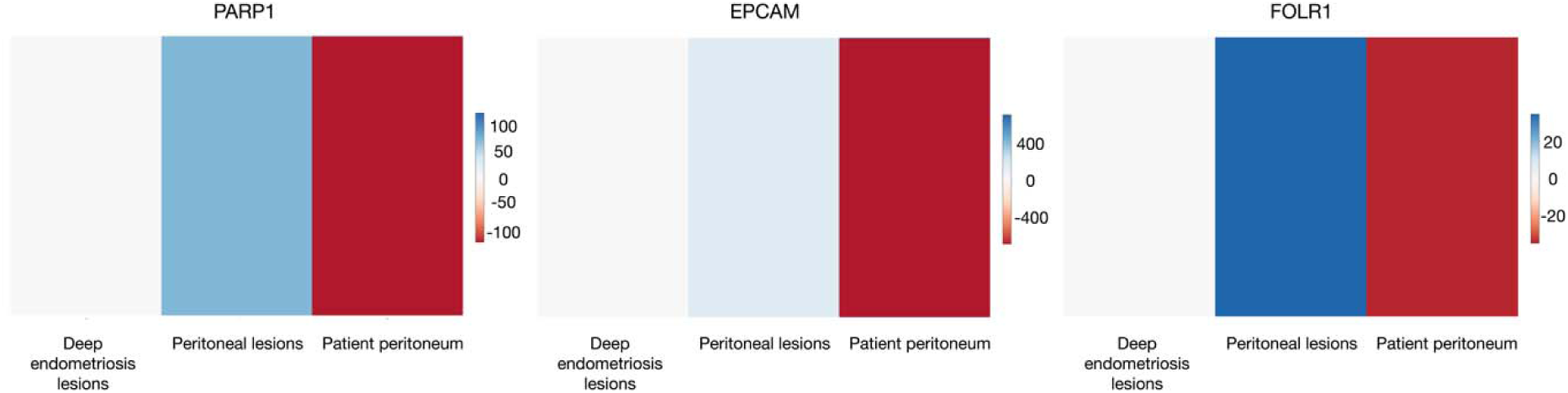
Heatmap depicting the expression of selected genes in control and endometriosis patient samples. Endometriotic lesions have a considerable increase in gene expression compared to the peritoneum. EndometDB is used to collect data.

### Immunohistochemistry analysis shows that PARP-1, EpCAM, and FRα are overexpressed in the endometriotic lesion compared to the surrounding tissue

Immunostaining for PARP-1 showed overexpression in foci of endometriosis relative to the surrounding tissue. Indeed, PARP-1 is mainly expressed by glandular epithelial cells and a few cytogenic stromal cells, with nuclear labeling ranging from moderate to strong degree of intensity. In the surrounding tissue, we observed a mild nuclear expression by stromal cells, including fibroblasts and endothelial cells, and immune elements with lymphocyte and macrophage morphology (**Figure 2**).

**Figure 2.**
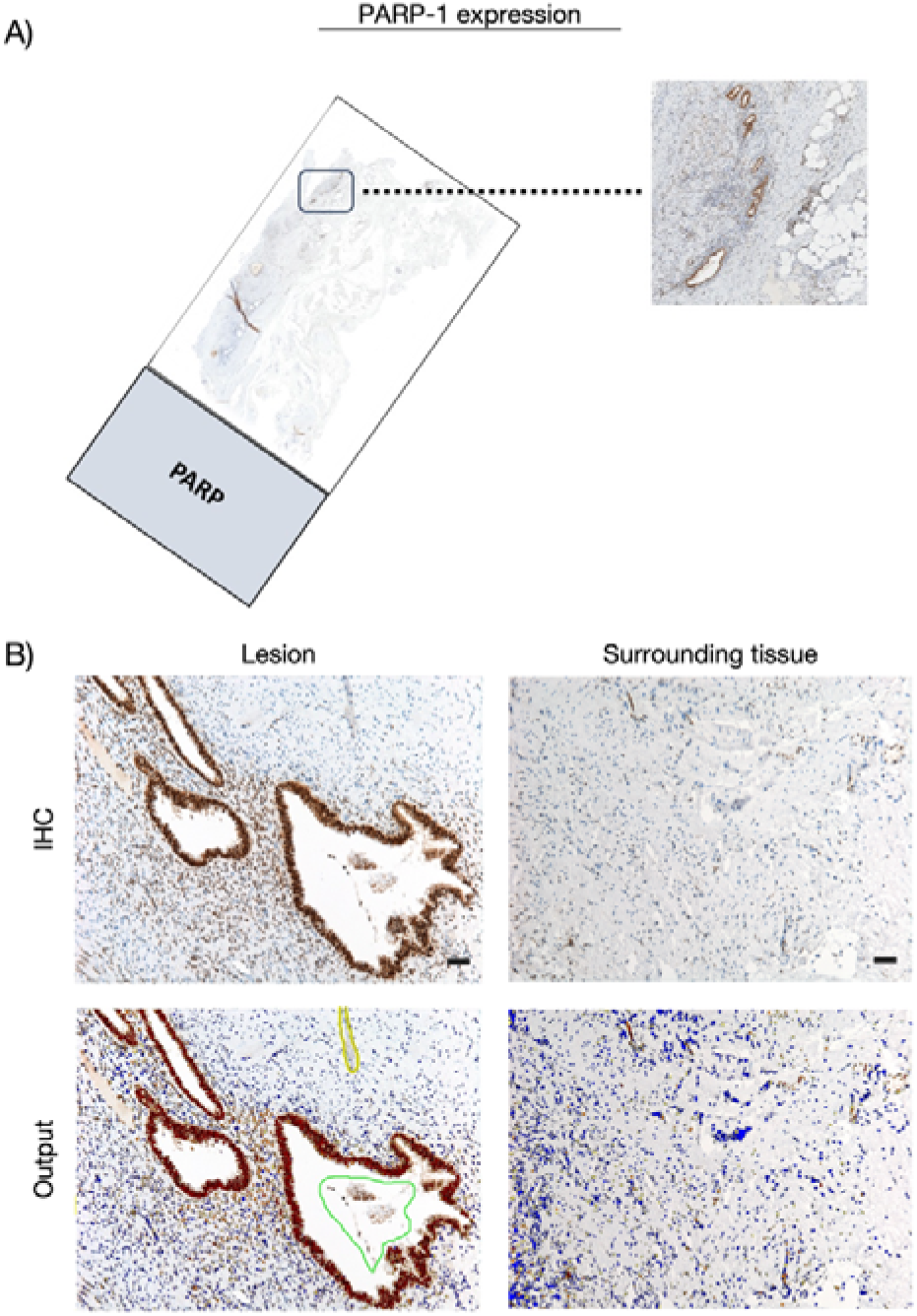
PARP-1 expression in the cohort of patients. **(A)** Whole slide image with a zoomed-in cross sectional area showing the endometriosis lesion. **(B)** Representative microphotographs showing expression of PARP-1 by IHC in endometriosis lesions and surrounding tissues. DAB (brown) chromogen was used to visualize the binding of the anti-human PARP-1 antibody. Magnification 100x; scale bars, 50 μm.

In endometriotic foci, with membrane and cytoplasmic labeling, immunohistochemical analysis revealed that glandular epithelial cells express EpCAM, exhibiting strong signal intensity **(Figure 3)**. Conversely, we observed few fibroblasts expressing EpCAM with membrane labeling and mild signal intensity in peri-endometriotic tissue.

**Figure 3.**
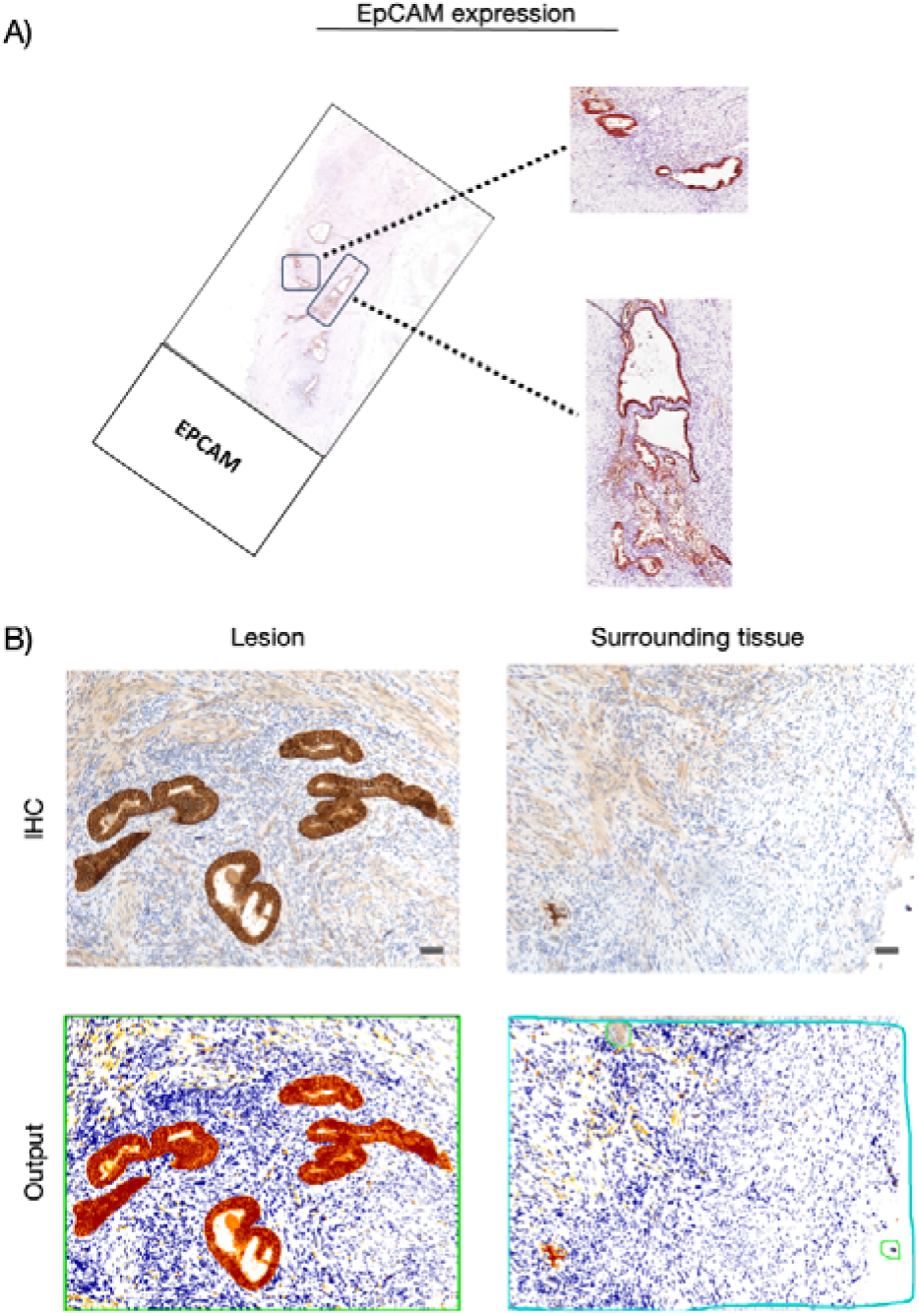
EpCAM expression in the cohort of patients. (**A**) Whole slide image with a zoomed-in cross sectional area showing the endometriosis lesion. (**B**) Representative microphotographs showing expression of EpCAM by IHC in endometriosis lesions and surrounding tissues. DAB (brown) chromogen was used to visualize the binding of the anti-human EpCAM antibody. Magnification: 100x; scale bars, 50 μm.

Regarding the FRα, we highlighted a higher expression in the endometrioid foci than the surrounding tissue **(Figure 4)**. Although with mild signal intensity compared to the other two markers, FRα is overexpressed in endometrioid glandular epithelial cells showing both membrane and cytoplasmic labeling. In contrast, surrounding tissue does not express it.

**Figure 4.**
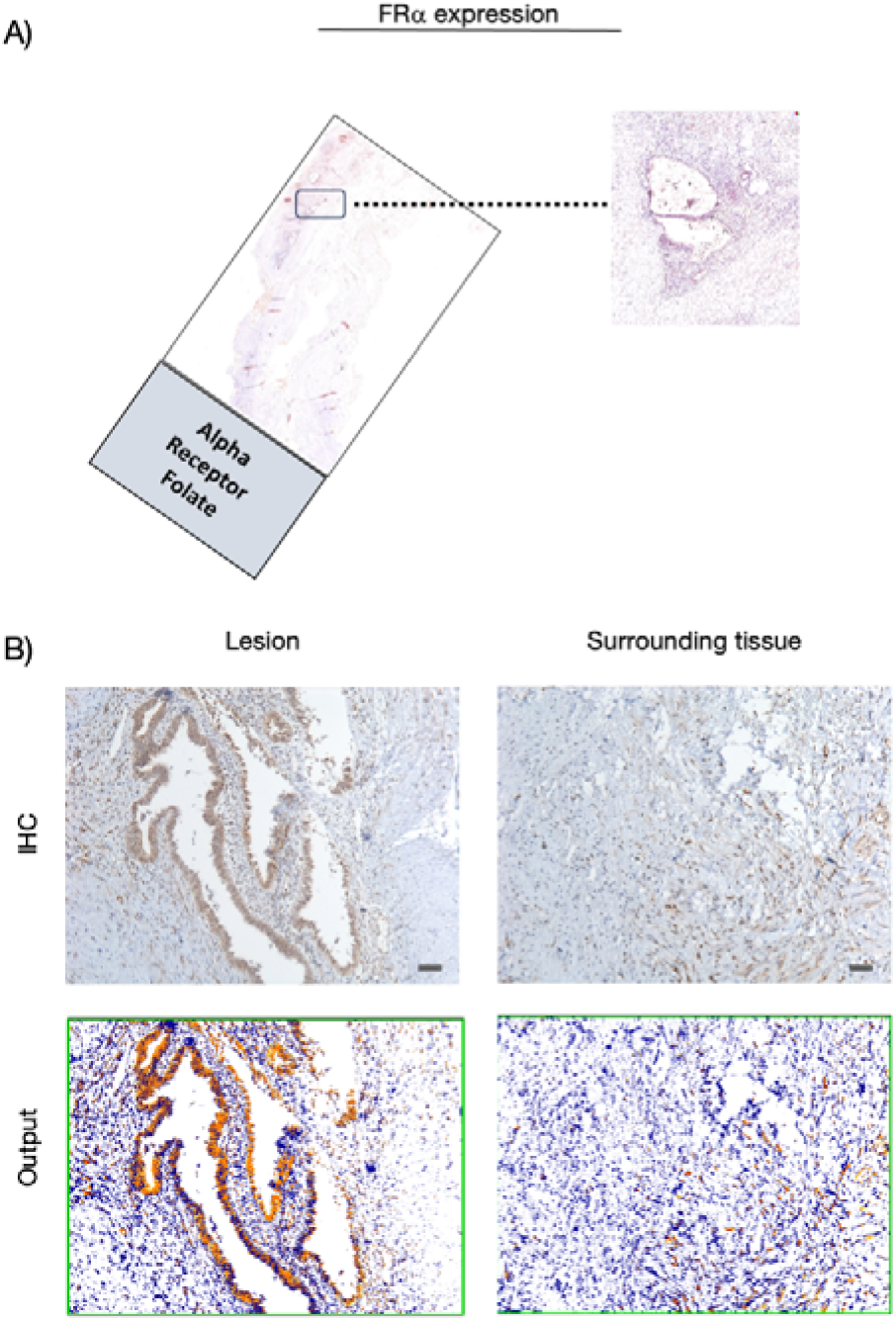
FRα expression in the cohort of patients. (**A**) Whole slide image with a zoomed-in cross sectional area showing the endometriosis lesion. (**B**) Representative microphotographs showing expression of FRα by IHC in endometriosis lesions and surrounding tissues. DAB (brown) chromogen was used to visualize the binding of the anti-human FRα antibody. Magnification 100x; scale bars, 50 μm.

Subsequently, this variable expression between endometriotic lesions and the surrounding area was confirmed by quantitative immunoassay analyses (**Figure 5**, **Additional file 1**, **2** and **3**).

**Figure 5.**
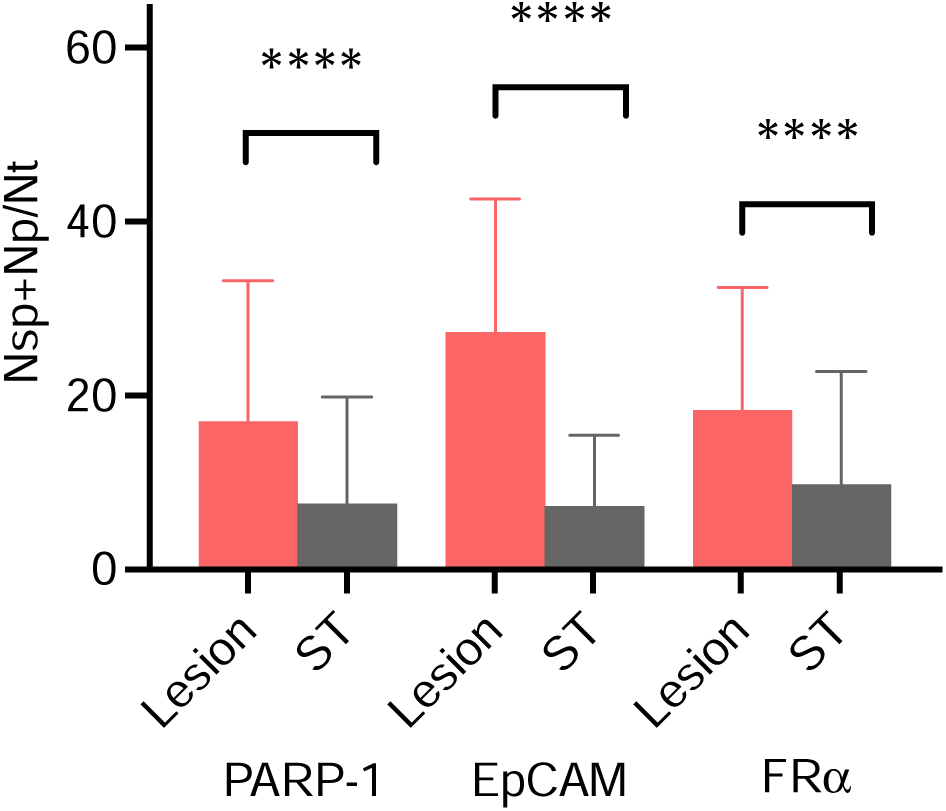
Comparison of the percentages of PARP-1, EpCAM, and FRα positivity between lesion samples and surrounding tissues (paired bootstrap t-test adjusted p-values < 10^-5^). ST: Surrounding tissue, Nsp: number of strong positive cells, Np: number of positive cells, Nt: number of total cells.

### Opal dual immunostaining shows a clear colocalization of PARP-1 with EpCAM and FRα within the endometriotic lesion

While investigating expression of PARP-1, EPCAM and FRα protein level in endometriosis foci by immunohistochemistry, we noticed that their expression increased almost the whole in endometriosis foci compared to surround tissue, independently by localization, as confirmed by quantification analyses.

Furthermore, we observed a considerable variability in terms of intensity signal among the considered markers and also heterogenous expression within the same marker.

Subsequently we carried out double immunofluorescence opal assays for PARP-1 and EPCAM and for PARP-1 and FRα. We consistently observed their expression in endometriosis foci, highlighting an intense PARP-1 nuclear expression, a strong EPCAM expression with membrane labeling and mild intensity of signal was detected for FRα. These data suggest that for intraoperative detection of endometriosis foci it might be useful the simultaneous employment of the two fluorescent probes in order to improve the quality and the radicality of surgery cure (**Figure 6**).

**Figure 6.**
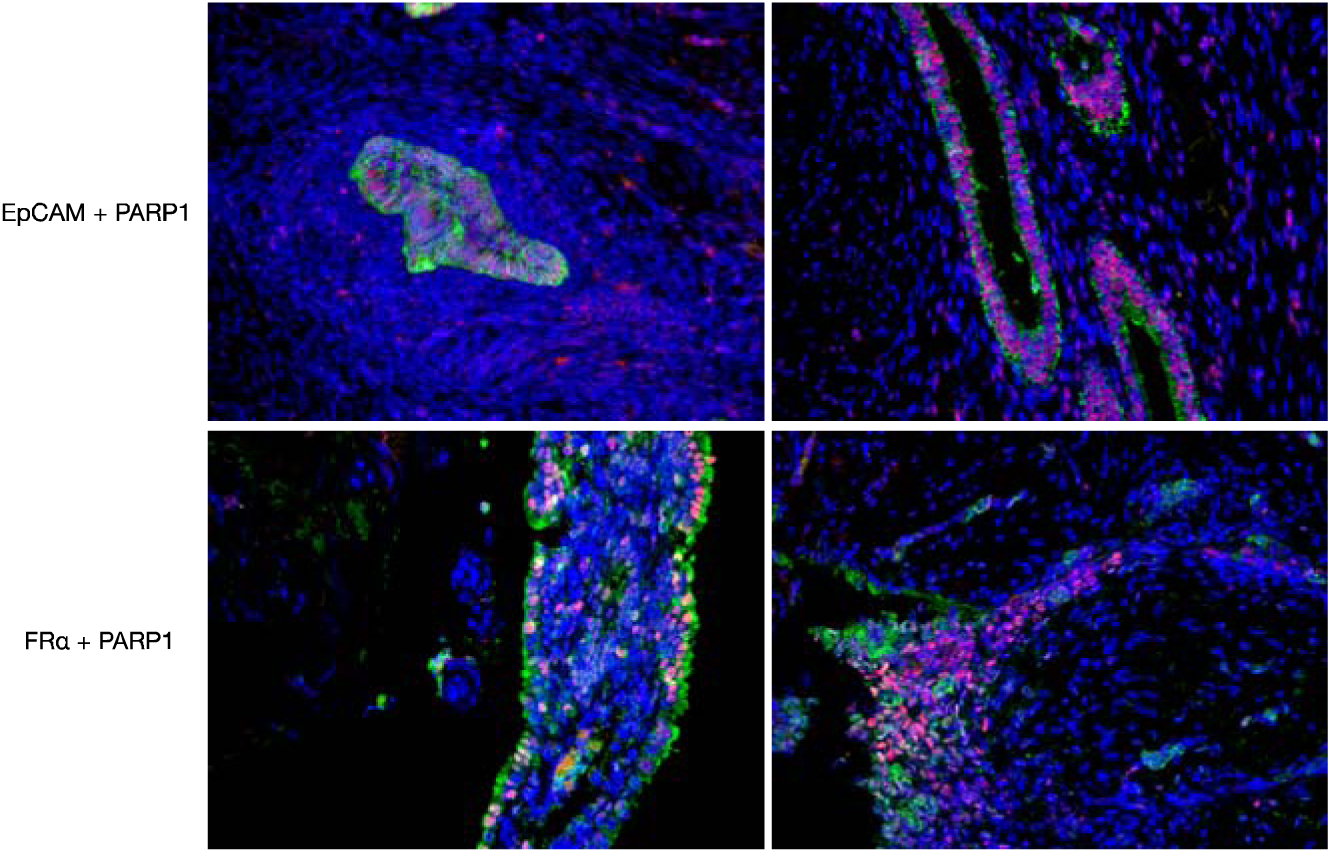
Representative microphotographs showing double immunofluorescence for EpCAM (green signal) and PARP-1 (red signal) and FRα (signal) and PARP-1 of endometriosis foci; nuclei were stained with DAPI. Magnification 200x).

## Discussion

The success of surgical excision in managing endometriosis holds great significance in improving patient outcomes. However, the identification of specific lesions can be challenging due to their small size or hidden location. To address this issue, image-guided surgery has emerged as a promising and innovative approach. This study aimed to evaluate the expression of three proteins that could potentially serve as targets for intraoperative endometriosis imaging. The proteins examined were EpCAM and FRα, both membrane proteins, and the nuclear marker PARP-1. The focus of the study was on endometriotic lesions that would benefit the most from a targeted probe imaging technique, excluding ovarian lesions that are easily visible. Instead, superficial and deep infiltrating endometriosis lesions were analyzed.

The significance of PARP-1 lies in its role as a biomarker for targeted therapy, as it is known to be overexpressed in various cancer types. In the field of molecular imaging, PARP inhibitors are utilized as a model to develop specialized contrast agents [22].

Particularly, PET imaging of PARP-1 has demonstrated successful results in noninvasively measuring physiologic levels of PARP-1 in patients and monitoring the therapeutic response to PARP inhibitor treatment [23]. Radiotracers are administered at low concentrations (subnanomolar) to minimize their pharmacological impact on the normal function of the PARP-1 enzyme. Moreover, the use of PARPi-FL has shown potential in assisting surgeons in detecting oral and tongue cancer, as well as determining the precise location and extent of the malignancy [24].

The significance of PARP-1 extends beyond its role in cancer, as it influences signaling pathways involved in immune response and inflammation. This suggests a potential rationale for developing radioligands to enable nuclear imaging of PARP-1 expression and activity in non-cancerous diseases such as cardiovascular disease, diabetes, and neurologic diseases [22]. This approach also aligns with the pathogenesis hypothesis of endometriosis, offering a new avenue for studying both PARP-1 and endometriosis [25,26]. However, there is currently a lack of data on PARP-1 expression in endometriotic lesions. A study by Barreta *et al.* examined PARP-1 immunohistochemistry expression in both carcinomas and endometriosis and revealed that benign ovarian lesions associated with endometriosis exhibit similar levels of PARP-1 expression as ovarian carcinomas linked to endometriosis [27]. These findings prompted our investigation into the expression of this marker in endometriosis, recognizing the importance of filling this knowledge gap.

Previous studies have provided evidence of PARP-1 expression in the uterus during crucial reproductive processes such as embryo implantation and decidualization, with its regulation being influenced by ovarian hormones [28]. These findings strongly indicate the involvement of PARP-1 in the intricate process of embryo implantation [28]. Imamura *et al.* further contributed to this knowledge by reporting on the role of PARPs in the pre-implantation development and epigenetic modification of mouse zygotes [29]. Additionally, the research conducted by Ménissier de Murcia *et al.* confirmed the significance of PARP-1 and PARP-2 in embryogenesis. Collectively, these studies emphasize the importance of PARP-1 in embryogenesis and highlight its potential as a target for further exploration in the field of reproductive medicine [30].

In our study, first we performed immunohistochemical analysis of the endometrium PPFE samples in both the proliferative and secretory phases, revealing the expression of PARP-1 in various components including endometrial glandular elements, cytogenic stroma elements, and uncommon immune elements. Interestingly, the secretory phase exhibited a more prominent expression profile, particularly in the stromal layer (**Additional file 4**). Further examination of endometriotic lesions confirmed the staining pattern observed in the endometrium, consistently allowing for the identification of lesion boundaries based on PARP-1 expression. Qualitative analysis of the lesions within their tissue environment revealed a notable contrast highlighted by PARP-1 expression, with lower levels of PARP-1 expression in the surrounding tissues and higher levels observed in areas with a stronger immune response. Using advanced image analysis algorithms, we confirmed that the number of PARP-1-positive nuclei was higher within the lesions compared to the surrounding tissues in the patients studied. These findings emphasize the significance of PARP-1 expression as a potential marker for distinguishing endometriotic lesions from the surrounding tissue, providing valuable insights into the pathophysiology and detection of endometriosis.

EpCAM expression was observed to be low in the surrounding tissues, highlighting a potential significant contrast between the background and endometriotic lesions, where its expression was high. However, relying solely on detecting and imaging the epithelial component of the lesions may be limited due to the histological variations and diverse proportions of stromal and epithelial cells within endometriotic lesions. The conventional histological assessment of a "typical" endometriotic lesion typically emphasizes the presence of endometrioid glands and/or stroma exclusively in ectopic sites [1,31,32]. Nonetheless, there are a few exceptions observed, such as stromal-only endometriosis or cases where the stroma is absent or replaced by adipocytes, histocytes, or inflammatory or fibrotic infiltrates, resulting in only identifiable glands being present [31,33,34]. These variations in histological composition underscore the complexity of endometriotic lesions, emphasizing the importance of comprehensive characterization methods beyond solely targeting the epithelial component. Despite extensive scientific papers and global research priority reports, it is evident that endometriotic lesions exhibit considerable variability in their appearance [35–37].

Recent studies have shed light on the histological heterogeneity of these lesions, which can vary not only across different individuals but also within the same individual and even within a single biopsy [38]. Recognizing this diversity, multiplexed imaging has emerged as an intriguing approach for gathering comprehensive information from patient tissue samples by simultaneously examining multiple biomarkers [39]. This technique involves combining targeted or untargeted dyes to address the tissue and anatomical structural heterogeneity observed in endometriosis. Multi-wavelength fluorescence imaging methods using a wide range of dye-functionalized targeting agents have been explored in preclinical animal models, revealing the potential of this approach [39]. Therefore, it is plausible to consider the utilization of a combination of epithelium- and stroma-specific probes to enhance the imaging and characterization of endometriotic lesions, enabling a more comprehensive understanding of their complex nature.

## Conclusions

Overall, the three markers exhibit considerable potential for the successful visualization of endometriosis. The findings of this study underscore the significance of PARP-1 expression as a potential biomarker for differentiating endometriotic lesions from surrounding tissue. With the anticipation of our results serving as a catalyst, we hope to inspire further investigations and the implementation of molecular imaging techniques in the pursuit of achieving precision surgery. By focusing on improving patient outcomes, the integration of molecular imaging has the potential to revolutionize the field, providing a more tailored and effective approach to addressing endometriosis. The potential of PARP-1 as a biomarker for endometriosis presents promising opportunities for additional research in understanding the underlying mechanisms of the disease and developing improved diagnostic and therapeutic strategies.

## Funding

This work was supported by the Italian Ministry of Health, through the contribution given to the Institute for Maternal and Child Health IRCCS Burlo Garofolo, Trieste, Italy.

## Disclosure

Disclosure: "none’. The authors declare no competing interests.

## Data availability

The histology images supporting the findings of this study are available in the open dissemination research data repository Zenodo with the digital object identifier (DOI): 10.5281/zenodo.8141561.

## Acknowledgements

Not applicable.

---

**Additional file 1**. Percentage of strong and weak PARP-1 positive cells quantified by the Nuclear v9 algorithm software in the lesion and surrounding tissues over focus. Weak positivity: signal intensity threshold 210; moderate positivity: signal intensity threshold 188; strong positivity: signal intensity threshold 162.

**Additional file 2**. Number of total cells (Ntot), number of EpCAM positive cells (Np), and number of EpCAM strong positive cells (Nsp) quantified by the Positive Pixel Count v9 algorithm (1+ weak positivity: signal intensity range 197–175; 2+ moderate positivity: signal intensity range 175–100; 3+ strong positivity: signal intensity range 100–0) in the lesion and surrounding tissues over focus.

**Additional file 3**. Number of total cells (Ntot), number of FRα positive cells (Np), and number of FRα strong positive cells (Nsp) quantified by the Positive Pixel Count v9 algorithm (1+ weak positivity: signal intensity range 197–175; 2+ moderate positivity: signal intensity range 175–100; 3+ strong positivity: signal intensity range 100–0) in the lesion and surrounding tissues over focus.

**Additional file 4**. Immunohistochemical analysis of the endometrium PPFE samples in both the proliferative and secretory phases.

## Notes

### Competing Interest Statement

The authors have declared no competing interest.

