## Additional file 4 for "PARP-1, EpCAM, and FRα as potential targets for intraoperative detection and delineation of endometriosis: a quantitative tissue expression analysis"

PARP-1 (clone EPR18461, Abcam ab19217, 1:1000 ph6)

Proliferative endometrium

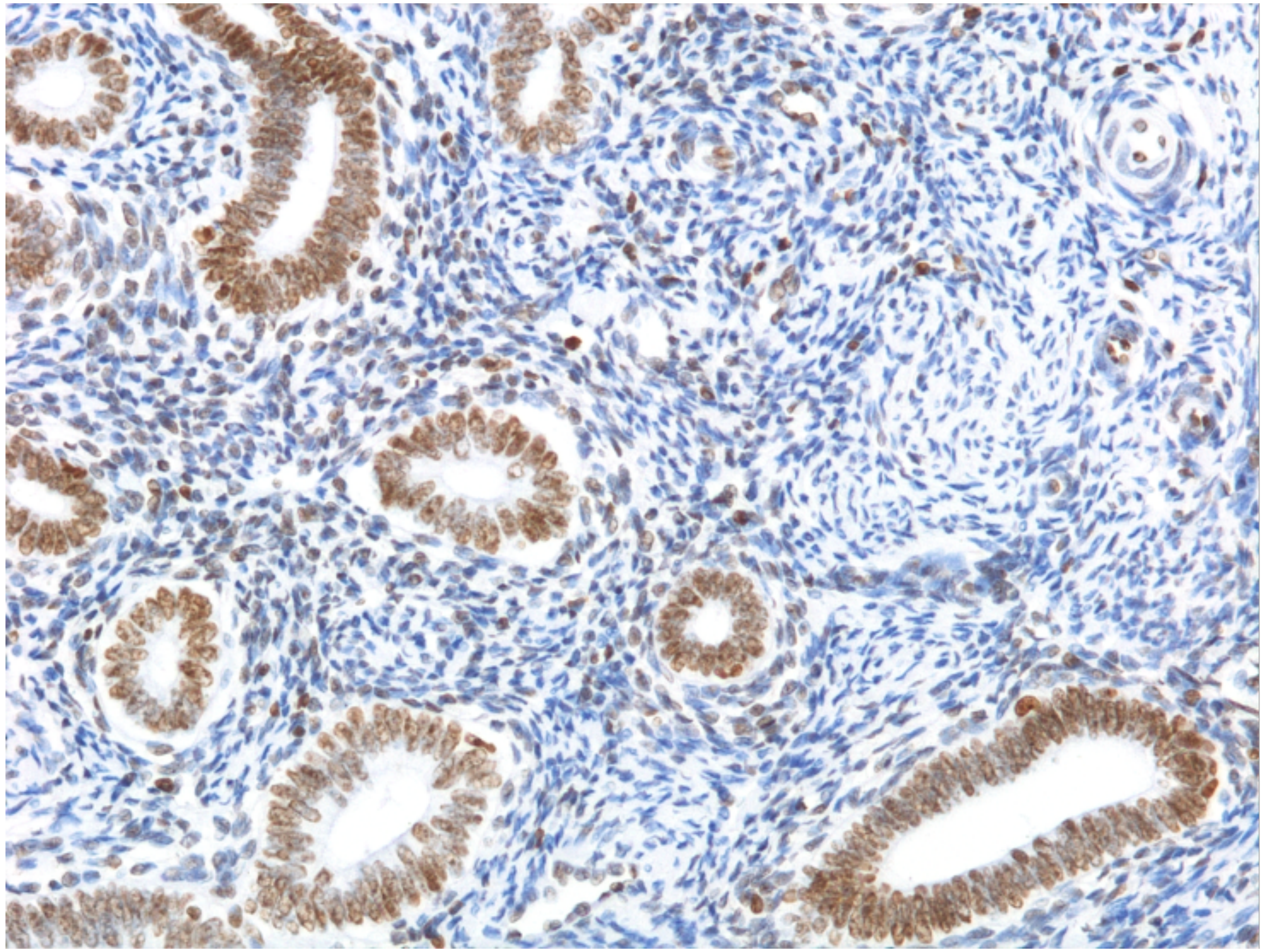

Secretory endometrium

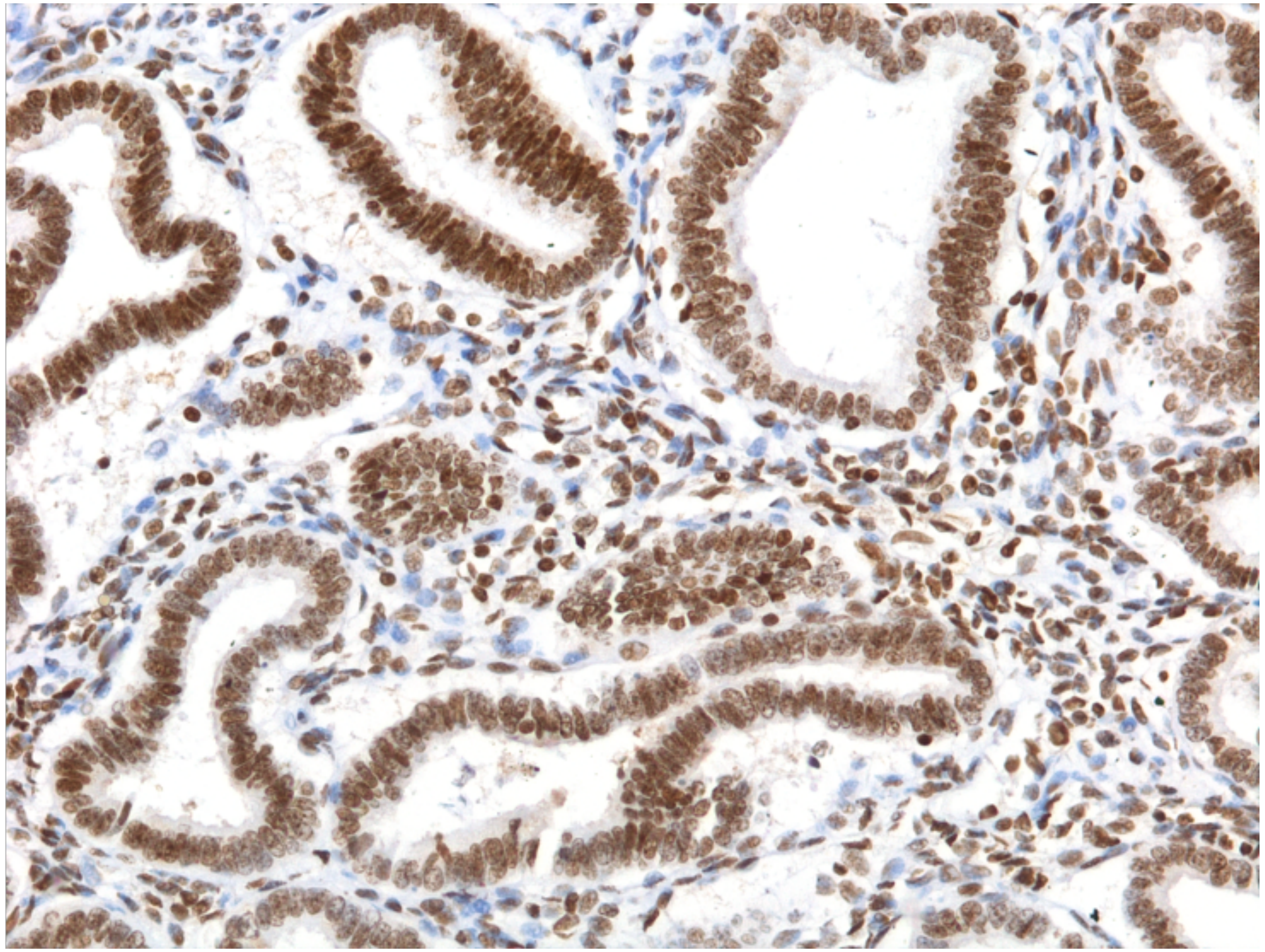

PARP-1 is expressed in various components, including endometrial glandular elements, cytogenic stromal elements, and uncommon immune elements. Interestingly, the secretory phase exhibits a more prominent expression profile, particularly in the stromal layer.
